# The effects of reference panel perturbations on the accuracy of genotype imputation

**DOI:** 10.1101/2023.08.10.552684

**Authors:** Jeremiah H. Li, Andrew Liu, C. Alex Buerkle, William Palmer, Gillian M. Belbin, Mohammad Ahangari, Matthew J.S. Gibson, Lex Flagel

## Abstract

Reference-based genotype imputation is a standard technique that has become increasingly popular in large-scale studies involving genomic data. The two key elements involved in the process of genotype imputation are (1) the haplotype reference panel to which a target individual is being imputed, and (2) the imputation algorithm used to infer missing genotypes in the target individual. The imputation literature has historically focused mainly on (2), with a typical comparative study investigating the relative performance of various imputation algorithms while holding the reference panel constant. However, the role of the reference panel itself (1) on overall imputation performance is equally, if not more, important than the choice among many high-performing algorithms. Even though it is intuitive that the quality of a reference panel should play a role in the accuracy of imputation, it is nonetheless unclear to what extent common errors during panel creation (e.g., genotyping and phase error) lead to suboptimal imputation performance. In this study, we investigate the effects of applying three distinct modes of perturbations to a widely used haplotype reference panel in human genetics on the resulting imputation accuracy. Specifically, we perturb the reference panel by (1) randomly introducing phase errors, (2) randomly introducing genotype errors, and (3) randomly pruning variants from the panel (all at varying magnitudes). We then impute a set of diverse individuals at various sequencing coverages (0.5x, 1.0x, and 2.0x) to these various perturbed panels and evaluate imputation accuracy using the *r*^2^ metric for the entire cohort as well as ancestry-stratified subsets. We observe that both phase- and genotype-perturbations can dramatically affect imputation accuracy, particularly at very low allele frequencies, while pruning variants has a far smaller effect. We then empirically verified that our simulations reliably predict the impact of potential filtering techniques in a real-world dataset. In the context of haplotype reference panels, these results suggest that phasing and genotyping accuracy are far more important than the density of a reference panel used for imputation.

## Introduction

Genotype imputation has become a standard element of the statistical geneticist’s toolkit in the last decade by “[enabling] researchers to inexpensively approximate whole-genome sequence data from genome-wide single-nucleotide polymorphism array data” Das et al. (2018). By allowing the reconstruction of haplotypes in a target individual at sites not directly assayed, imputation effects an increase in the statistical power of association studies, more accurate fine-mapping, and provides an elegant solution to integrating results from samples assayed on different platforms. More recently, reference-based imputation from low-coverage whole-genome sequence data (or low-pass sequencing) has emerged as an appealing alternative to imputation from genotyping arrays for a variety of reasons, including the lack of ascertainment bias, overall higher imputation accuracy, and the more flexible nature of sequence data as compared to fixed genotyping array calls (Li et al., 2021; Wasik et al., 2021; Rubinacci et al., 2021, 2023; Li et al., 2022, 2011).

The typical setup for genotype imputation from whatever data modality (whether it be genotyping array calls or raw sequence data) for a given cohort requires the selection of two key elements — the haplotype reference panel with which a target individual is to be imputed, and the imputation algorithm itself. Most modern imputation algorithms are based on the Li & Stephens haplotype copying model, introduced by the eponymous authors in Li and Stephens (2003), wherein a target individual’s underlying haplotypes are modeled as an imperfect mosaic of the haplotypes in the reference panel via the use of a hidden markov model (HMM). The last decade has seen advances in the algorithmic methods used for imputation, with the most recent algorithms allowing whole-genome imputation in humans from low-pass sequence data at a computational cost of *£*0.08 per sample (Rubinacci et al., 2023).

Indeed, as of 2022, there exist at least a half-dozen viable (in terms of computational tractability and cost) methods for genome-wide imputation; the runtime, accuracy, and ease of use of these methods far outstrip the original implementations of the Li & Stephens model for this purpose (Das et al., 2018).

As a result of the diverse selection of extant methods, there exists a rich literature of comparative studies in which imputation algorithms are pit head-to-head and their relative performance evaluated. For several examples, see Hui et al. (2020), De Marino et al. (2022), Roshyara et al. (2016), and Pook et al. (2020). Similarly, it is also common practice for any study introducing a new imputation method to evaluate its performance against existing methods by imputing the same set of target individuals to a given reference panel while varying the imputation algorithm used (Browning and Browning, 2016; Rubinacci et al., 2021; Browning et al., 2018; Das et al., 2016).

The other crucial element of any imputation pipeline is the haplotype reference panel itself; *i.e*., the set of source haplotypes from which the Li & Stephens model HMM draws to model the target individual’s haplotypes. This takes the form of a complete and phased genotype matrix. The key elements of a reference panel that affect imputation accuracy are understood to be (1) its size (number of haplotypes), (2) the site frequency spectrum of the variants in the panel, and (3) the haplotype accuracy of its constituent samples (Das et al., 2018).

At a high level, most modern large haplotype reference panels are constructed as follows: (1) high-depth short-read whole genome sequence data is generated for a set of samples, from which (2) genotypes are (joint-) called genome-wide, (3) missing genotypes are filled in using within-cohort imputation methods such as `beagle4` (Browning and Browning, 2007), and (4) statistical phasing is performed to generate complete haplotypes from the non-missing genotype matrix.

Methods for statistical phasing have also undergone significant improvement within the last decade, with algorithms like `SHAPEIT5` and `beagle5` (Hofmeister et al., 2023; Browning et al., 2021) enabling rapid and accurate haplotype inference from genotypes alone. One typical metric of interest when comparing phasing algorithms on the basis of the accuracy of reconstructed haplotypes is the so-called “switch error rate” (SER), which is the proportion of consecutive heterozygotes that are incorrectly phased (typically evaluated by comparison to gold-standard haplotypes generated using orthogonal technologies such as long read sequencing) (Choi et al., 2018). Similarly to the literature on imputation methods, the typical study comparing statistical phasing methods focuses on the resulting differences in SERs when a variety of methods are applied to the same dataset.

Given the above, it is clear that the quality of a haplotype reference panel in terms of haplotype accuracy (*i.e*., phasing and genotype accuracy) *should* be of paramount importance in the process of panel construction. However, there does not appear to be a direct evaluation of the effects of phasing and genotyping errors in a reference panel on downstream imputation accuracy, which is required to give a sense of how important these various factors are in practice. This information would be valuable in study design and evaluation of the return on improving a reference panel through larger numbers of sites and greater haplotype accuracy.

Here, we take a subset (chromosome 20) of a haplotype reference panel widely used in human genetics, the New York Genome Center’s high-depth resequencing effort of the 1000 Genomes Phase 3 (plus additional related individuals) (which we denote as the “NYGC dataset”) (1000 Genomes Project Consortium, 2015; Byrska-Bishop et al., 2022), and apply to it three different types of perturbations at various magnitudes:

1. Phase perturbations: at a set of randomly selected heterozygotes, flip the phase (*i.e*., 0|1 *→* 1|0 or 1|0 *→* 0|1).
2. Genotype perturbations: at a set of randomly selected alleles, flip the genotype (*i.e*., 0 *→* 1 or 1 *→* 0) while preserving phase.
3. Density perturbation: prune the panel of a randomly selected set of variants.

The first two perturbation types are designed to investigate the direct effects of increasing levels of phasing and genotyping errors on downstream imputation accuracy, whereas the third is designed to investigate the direct effects of the density of sites in the reference panel. Though panel density is largely determined by the number of individuals comprising the panel, various filtering procedures and postprocessing steps applied to any particular cohort during the process of panel creation can result in substantially different densities in the resulting panel. For instance, the 1000 Genomes Phase 3 reference panel comprising 2504 individuals contains *∼*88M variants, whereas the Haplotype Reference Consortium panel comprising 64, 976 individuals contains *∼*39M variants (1000 Genomes Project Consortium, 2015; Haplotype Reference Consortium, 2016).

Following the creation of these perturbed panels, we selected a group of 60 diverse individuals from the original 1000 Genomes Phase 3 (1KGP3) project, downsampled them to the equivalent of 0.5x, 1.0x, and 2.0x coverage, and imputed them using `GLIMPSE v1.1.1` (Rubinacci et al., 2021) to the perturbed panels as well as the original, unperturbed panel. We then evaluated imputation accuracy for the entire cohort as well as ancestry-stratified subsets of the cohort by means of the imputation *r*^2^ metric, which has a direct interpretation with the statistical power of an association test for a binary trait at a given variant (Das et al., 2018; Pritchard and Przeworski, 2001).

We find that while both phase and genotype perturbations can significantly affect the accuracy of imputation – particularly in the low allele frequency domain – pruning the panel of variants, even when 50% of variants are removed, has a relatively small effect on overall imputation accuracy. To provide an empirical test of the directional effect of reducing genotyping error, we then filtered out *≈*10% of the unperturbed panel based on a genotype quality filter to achieve a callset with theoretically more accurate genotypes and repeated the *r*^2^ analyses. We found that at all coverages, this filtered panel demonstrated an improved *r*^2^ with a particularly pronounced impact at MAFs ≲ 1%.

## Results

### Experimental design

#### Low-coverage sequence data

To compare performance across a range of diverse ancestries, we selected twelve individuals from each superpopulation group in the 1000 Genomes Phase 3 release (AFR, AMR, EAS, EUR, SAS) for a total of 60 individuals under analysis. Since the NYGC’s release of the 1KGP3 dataset includes 602 trios, which could enable higher imputation accuracy for those samples if related individuals were not excluded, we specifically chose individuals that were not part of those trios.

For each individual, we downloaded the high-depth sequence data used for creation of the NYGC haplotype reference panel from the International Genome Sample Resource website and downsampled the reads to 0.5x, 1.0x, and 2.0x coverage to simulate low-pass sequence data. We then took the downsampled reads that aligned to chromosome 20 of GRCh38, generated genotype likelihoods using a custom tool, and imputed them up to the original, unperturbed NYGC reference panel in addition to all perturbed panels in a leave-one-out manner (Methods).

#### Reference panel perturbations

As mentioned above, we selected the NYGC dataset, a modern haplotype reference panel deriving from high-coverage (30x) short-read Illumina sequences (Byrska-Bishop et al., 2022). For simplicity, we limited all analyses to a reduced version of the reference panel comprising only bi-allelic SNPs on chromosome 20 with all singletons removed. The resulting reference panel comprises a total of 1, 391, 020 polymorphic sites.

We then created a total of 17 perturbed (eight phase-perturbed, four genotype-perturbed, and five pruned) reference panels for a total of 18 panels including the original, unperturbed panel. The range of perturbation magnitudes were chosen (1) to investigate the robustness of imputation to extreme phase misspecification, (2) to probe a range of realistic genotyping error rates, and (3) to span a range of high densities found in modern haplotype callsets.

To create each of the phase-perturbed panels, we introduced random phase errors at heterozygotes in the original panel by walking along the entire genotype matrix and drawing from a Bernoulli distribution Bernoulli(*p*) at each heterozygote, flipping the phase (*i.e*., 0|1 *→* 1|0 or 1|0 *→* 0|1) if the draw was successful. To encompass a range of phase errors, we created a panel for each *p ∈ {*0.001, 0.01, 0.05, 0.1, 0.2, 0.3, 0.4, 0.5*}*, such that for *p* = 0.5, *≈*50% (up to the randomness in the draws) of the original heterozygotes have opposite phase compared to the original panel.

To create each of the genotype-perturbed panels, we introduced random genotyping errors into the original panel by walking along the entire genotype matrix and drawing from a Bernoulli distribution Bernoulli(*p*) at each *allele* (*i.e*., for a genotype call 1|0, we draw from the distribution twice; once for the first allele 1 and once for the second allele 0). If the draw was successful, we flip the allele from the reference to alternate allele or vice versa (*e.g*., if the original genotype was 1|0 and we have a successful draw at the first but not second allele, the resulting perturbed genotype is 0|0). We varied *p* to create a panel for each *p ∈ {*0.001, 0.005, 0.01, 0.05*}* to model a range of genotyping error, for a total of five genotype-perturbed reference panels.

To create each of the pruned panels, we randomly selected a proportion *p* of the variants in the original reference panel and removed them entirely from the reference panel. We varied *p* to create a panel for each *p ∈ {*0.1, 0.2, 0.3, 0.4, 0.5*}* to encompass a range of resulting panel densities.

We then imputed the same set of sequence reads at the various downsampled coverages up to the density of these eighteen resulting reference panels. To evaluate imputation performance, we use the imputation *r*^2^ metric, which provides a direct interpretation in terms of statistical power and sample size for association tests for binary traits (Methods).

### Overall results

For each downsampled cohort, we imputed all samples to each of 18 perturbed panels, for a total of ((3 coverages) *×* (60 individuals) *×* (18 panels) = 3240) sample runs. This section focuses on the 1.0x coverage data to illustrate quantitative trends. For all modes of perturbation, the original panel (no perturbation) performed the best, with decreasing accuracy across the MAF spectrum with increasing magnitude of perturbation ^1^.

*∼*

Interestingly, we observe that pruning the panel produces a much smaller change in accuracy compared to the other two types of perturbations, even at rare variants; indeed, at the lowest MAF bin ([0.0001, 0.0005]), pruning the panel by 50% only effects a decrease in mean *r*^2^ (Methods) from 0.287 to 0.241 (a 16% decrease) compared to the unperturbed panel for samples at 1.0x. On the other hand, even small perturbations to the genotypes cause dramatic decreases in imputation accuracy; genotyping error at a rate of 0.1% effects a threefold decrease in mean *r*^2^ from 0.287 to 0.0920 (a 68% decrease) at 1.0x. Similarly, perturbations to the phase at about *∼*5% begin to seriously affect the accuracy at rare variants, with a near two-fold decrease in mean *r*^2^ from 0.287 to 0.128 (a 55.4% decrease) at the lowest MAF bin. This is seen at the genome-wide level as well, with a proportional decrease of 6.58% in genome-wide *r*^2^ for the maximally pruned panel and a proportional decrease of 21.1% for the 0.01% genotype-perturbed panel and 20% for the 5% phase-perturbed panel.

The effect of perturbations at all magnitudes are particularly amplified at the low end of the minor allele frequency spectrum, while imputation at common variants remains surprisingly robust regardless of the magnitude of perturbation. Indeed, all phase- and genotype-perturbed panels except for the 5%-genotype-perturbed panel still display imputation *r*^2^s of *>* 0.9 in the common variant (MAF *>* 0.1) regime. This result is particularly interesting to note given the magnitude of some of these perturbations – for instance, the maximally phase-perturbed panel corresponds to one in which a full 50% of heterozygotes have had their phase status modified — equivalent to completely random phasing.

However, perhaps the most interesting observation is that even small genotype perturbations significantly attenuate imputation accuracy, even at the smallest perturbation value of 0.001, or 0.1%. For instance, for the 1.0x sequencing data, the average genome-wide *r*^2^ decreased by 21% relative to the unperturbed panel.

This error rate is well within the range of genotyping errors seen in modern sequence callsets (Wall et al., 2014; Byrska-Bishop et al., 2022). Given previous observations in the literature that genotyping error rates typically increase with decreasing MAF and the observation made here that the effects of perturbations are also amplified with decreasing MAF, this result emphasizes the importance of accurate variant calls at rare variants (Wall et al., 2014).

### The effect of sequencing coverage

Intuitively, imputation performance should increase with higher sequencing coverages since increased coverage means that the HMM has more observed data with which to infer the true underlying haplotypes. We therefore examined the imputation performance of these perturbed panels at three different coverages within the range of what is typically considered “low-pass sequencing”: 0.5x, 1.0x, and 2.0x.

Figure 1 depicts imputation accuracy for the entire cohort at each coverage (columns) and perturbation type (rows). As expected, the imputation *r*^2^ for cohorts at a higher coverage is consistently higher than those with lower coverage.

**Figure 1:**
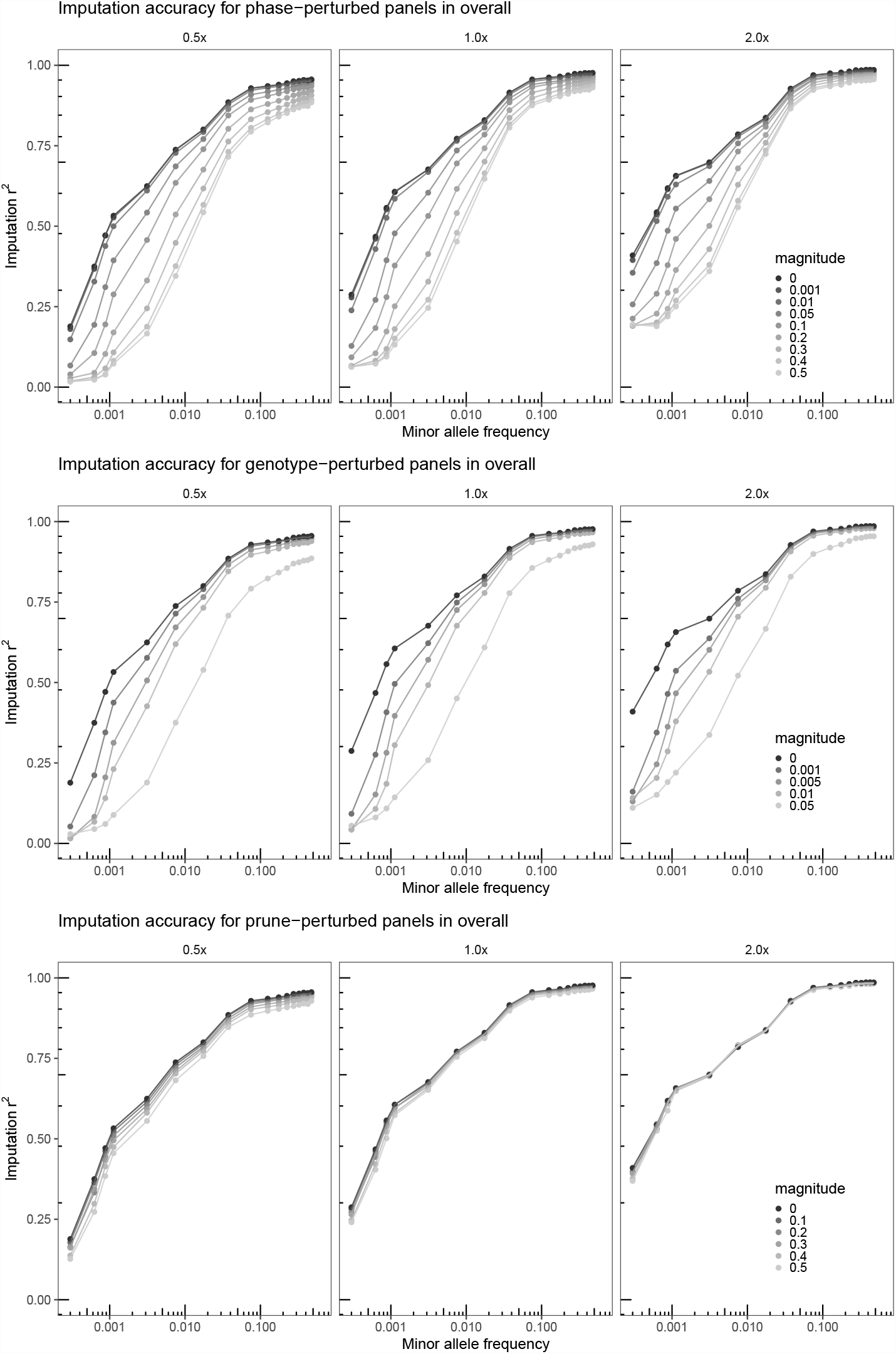
Imputation accuracy by minor allele frequency for sequence data at 0.5x, 1.0x, and 2.0x coverages. Each pane depicts imputation *r*^2^ for the full cohort of 60 individuals as evaluated against the unperturbed genotypes, and each line describes performance for the cohort for a particular panel. Note that the relative effects of the perturbations are diminished with increasing coverage particularly for rarer variants.

To quantitatively investigate the relative reduction in imputation accuracy induced by perturbations as a function of coverage, perturbation type, and magnitude, we computed the genome-wide average imputation *r*^2^ for a given cohort divided by the corresponding value for the unperturbed panel. Figure 2 depicts this quotient, and illustrates several important observations. There is an obvious trend toward lower relative performance with increasing perturbation at a fixed coverage, corresponding with our previous observations, but interestingly, the *rate* of decrease with respect to perturbation magnitude is magnified with decreasing coverage. Intuitively, if one thinks about the reference panel as a statistical prior and the input sequences as evidence with which to update the prior, then fewer observations correspond with fewer updates, and the effect of the prior is stronger (Stephens and Donnelly, 2003). In this context, this means that as the “prior” (reference panel) becomes increasingly misspecified (perturbed), the importance of the observations is increased, and thus the relative performance decrease is ameliorated at higher sequence coverages.

**Figure 2:**
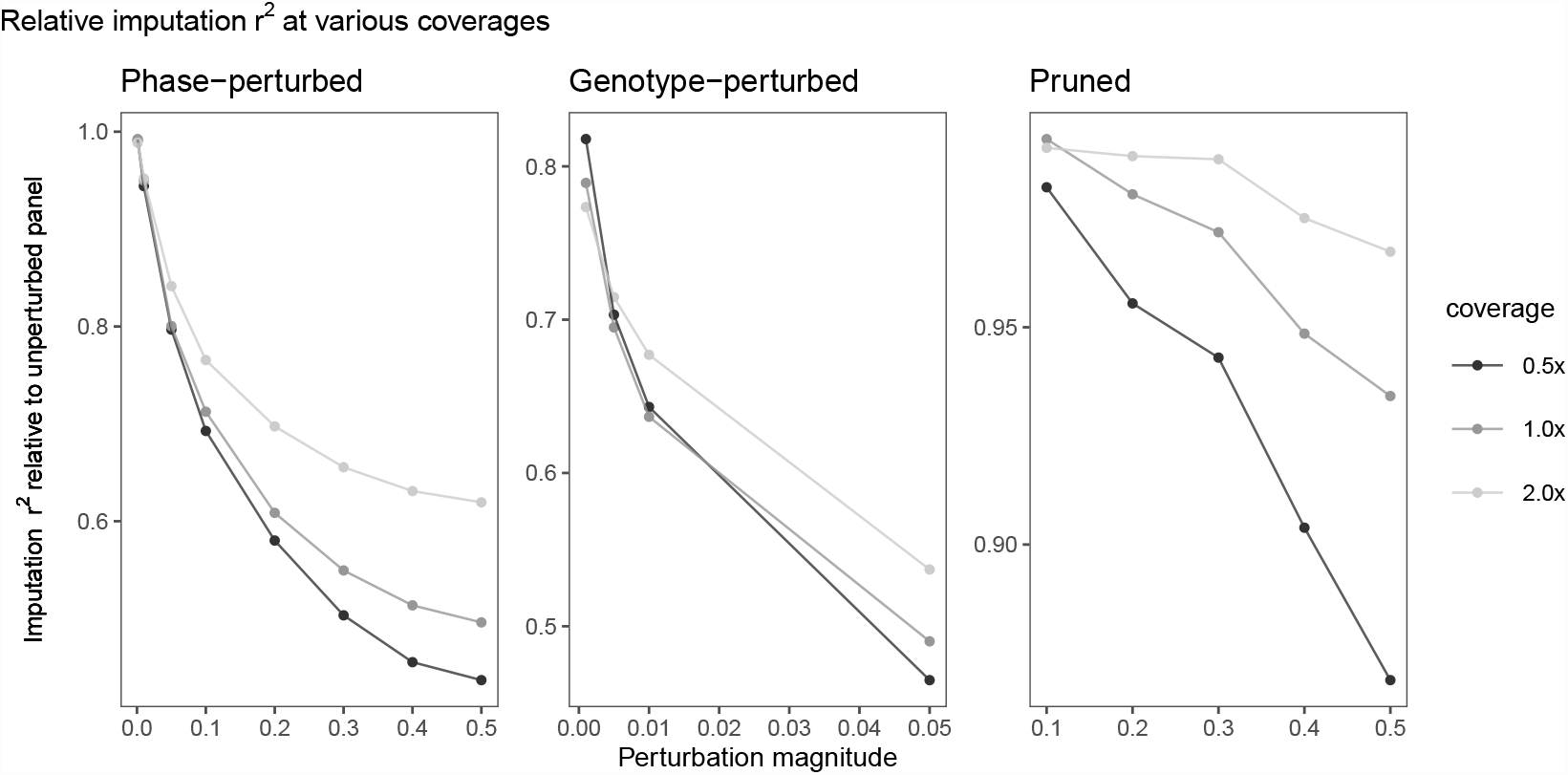
Relative imputation *r*^2^ for the entire cohort compared to the unperturbed panel. Each pane depicts a mode of perturbation and the y-axis depicts the cohort’s average *r*^2^ across all variants divided by the *r*^2^ corresponding with the unperturbed panel, and each line depicts a constant sequencing coverage. Note that the relative imputation accuracy decreases with the magnitude of perturbation and that on average, higher sequencing coverages retain greater proportional accuracy compared to lower coverages.

### The relationship between phase perturbation and switch error rate

The switch error rate of a set of phased haplotypes for a single individual is the “proportion of heterozygous sites that are phased incorrectly relative to the previous heterozygous site,” which measures the degree to which phase was incorrectly inferred, and serves as the primary metric of interest when evaluating the phased callset resulting from applying certain phasing algorithms or sequencing technologies (*e.g*., long-read sequencing by its nature allows much better phasing across covered regions) (Scheet and Stephens, 2006).

While our phase perturbation method of randomly flipping heterozygous sites is similar to inducing switch errors, there is the additional complication due to the fact that the switch error rate only concerns pairs of consecutive incompatibly phased heterozygotes rather than just the number of nominally incorrectly phased heterozygotes (in which case it would simply be the probability of success in the Bernoulli trial in the expectation).

However, we can empirically calculate the SER *induced* by the random perturbations, which provides a useful interpretation of the perturbations performed here in terms of a more familiar metric. Figure 3 depicts the relation between induced switch error rate and the magnitude of the phase perturbations, which appears approximately sigmoidal. Each violin plot describes the distribution of switch errors introduced calculated on a per-sample level, such that each describes a distribution across the 3202 individuals within the reference panel. The relation is unsurprising in the sense that greater random perturbation corresponds with greater induced switch error rates in a monotonic manner, with the minimal mean SER coming out at 0.2% and the maximal mean SER topping out at 50%.

**Figure 3:**
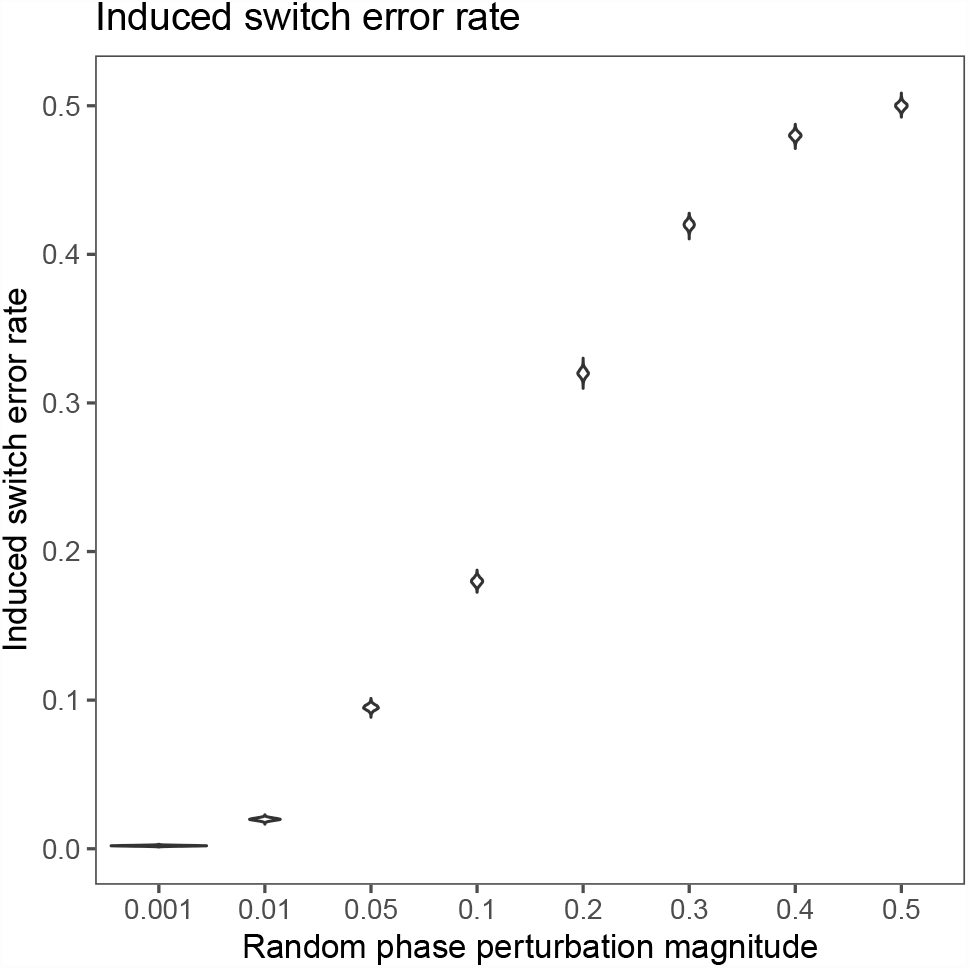
Switch error rate induced by random phasing perturbations. Here, the x axis is the magnitude of the phase perturbation, the y axis is the calculated switch error rate compared to the unperturbed callset, and each violin plot describes the distribution of switch error rates at an individual level. Thus, each violin plot describes the distribution of induced switch error rates for the 3202 individuals in the panel for each perturbation level.

While the primary purpose of this experiment is to investigate the robustness of imputation to high levels of phase misspecification, it is useful to place the values of the induced SERs in the context of modern phasing algorithms and genotyping technologies.

Historically, the first high-throughput statistical phasing methods were developed for data deriving from genotyping arrays, although the application to sequence-derived genotypes was quickly adopted. Methods such as `SHAPEIT`, `MaCH`, `Fastphase`, and `Beagle` applied to genotyping array data and low-coverage sequence data regularly achieved genome-wide SERs in the mid-single-digit percentage range, comparable to the lower end of the investigated perturbation spectrum (Browning and Browning, 2007; Delaneau et al., 2013; Williams et al., 2012). As a contrast, the highest end of the perturbation range (50%), corresponds to an algorithm that simply performs random haplotype phasing.

However, with the advent of more affordable whole genome sequencing, extremely large reference panels, and the ongoing refinement of phasing methodologies, modern methods applied to high-depth short-read WGS data regularly achieve SERs of well under 1%, which can be further reduced with the use of long-read sequencing technologies and hybrid methods such as the inclusion of parents to enable trio phasing and the augmentation of WGS data with Hi-C or Strand-seq data (Hofmeister et al., 2023; Byrska-Bishop et al., 2022; Browning et al., 2021; Choi et al., 2018; Lin et al., 2022). In this context, it is useful to look at the *r*^2^ attenuation for a realistic SER: for instance, the phase-perturbation with magnitude 0.001, which corresponds with a SER of 0.2%. We see that the effect is minimal (Figure 1), with a decrease of only 0.725% in genome-wide *r*^2^, indicating that results of modern phasing algorithms are sufficient for the purposes of creating reference panels used in imputation.

### Results across ancestries

The sample set chosen for this study represents global diversity to investigate the interaction of ancestry with imputation accuracy. Figure 4 illustrates the effect of ancestry (compare to Fig. 2 for overall 1.0x coverage performance in the superpopulation). We observe different slopes for different ancestries, which confirms differential effects of perturbation by subpopulation. Interestingly, there are several population/coverage/perturbation magnitude combinations in which the perturbed panels yield a slightly *higher* mean imputation *r*^2^ than the unperturbed panel, a pattern not observed when looking at the overall cohort (Figure 2). Supplementary Table 1 shows these cases.

**Table 1:** Table describing the genome-wide weighted aggregated mean *r*^2^ for various coverages. Here, “Unfiltered *r*^2^” refers to the *r*^2^ obtained by imputing to the original, unfiltered panel, “unfiltered *r*^2^ (intersect)” refers to the *r*^2^ of the same but evaluated only at the intersection of variants between the filtered and unfiltered panels, and “GQ-filtered *r*^2^” refers to the *r*^2^ obtained by imputing to the filtered panel. The last two columns are ratios of the *r*^2^s obtained from imputation for the filtered panel to the *r*^2^s obtained by imputing to the unfiltered panel at both all unfiltered sites and only at sites in the intersection. Note that imputing to the filtered results effects an improvement in *r*^2^ at both shared sites and sites retained after filtration.

| Coverage | Unfiltered $r^2$ | Unfiltered $r^2$<br>(intersect) | GQ-filtered $r^2$ | (GQ-filtered $r^2$)<br>/ (Unfiltered $r^2$<br>(intersect)) | (GQ-filtered $r^2$)<br>/ (Unfiltered $r^2$) |
| --- | --- | --- | --- | --- | --- |
| 0.5x | 0.4814 | 0.5135 | 0.5180 | 1.00879 | 1.0758 |
| 1x | 0.5517 | 0.5996 | 0.6078 | 1.01376 | 1.1017 |
| 2x | 0.6181 | 0.6917 | 0.6960 | 1.00618 | 1.1259 |

**Figure 4:**
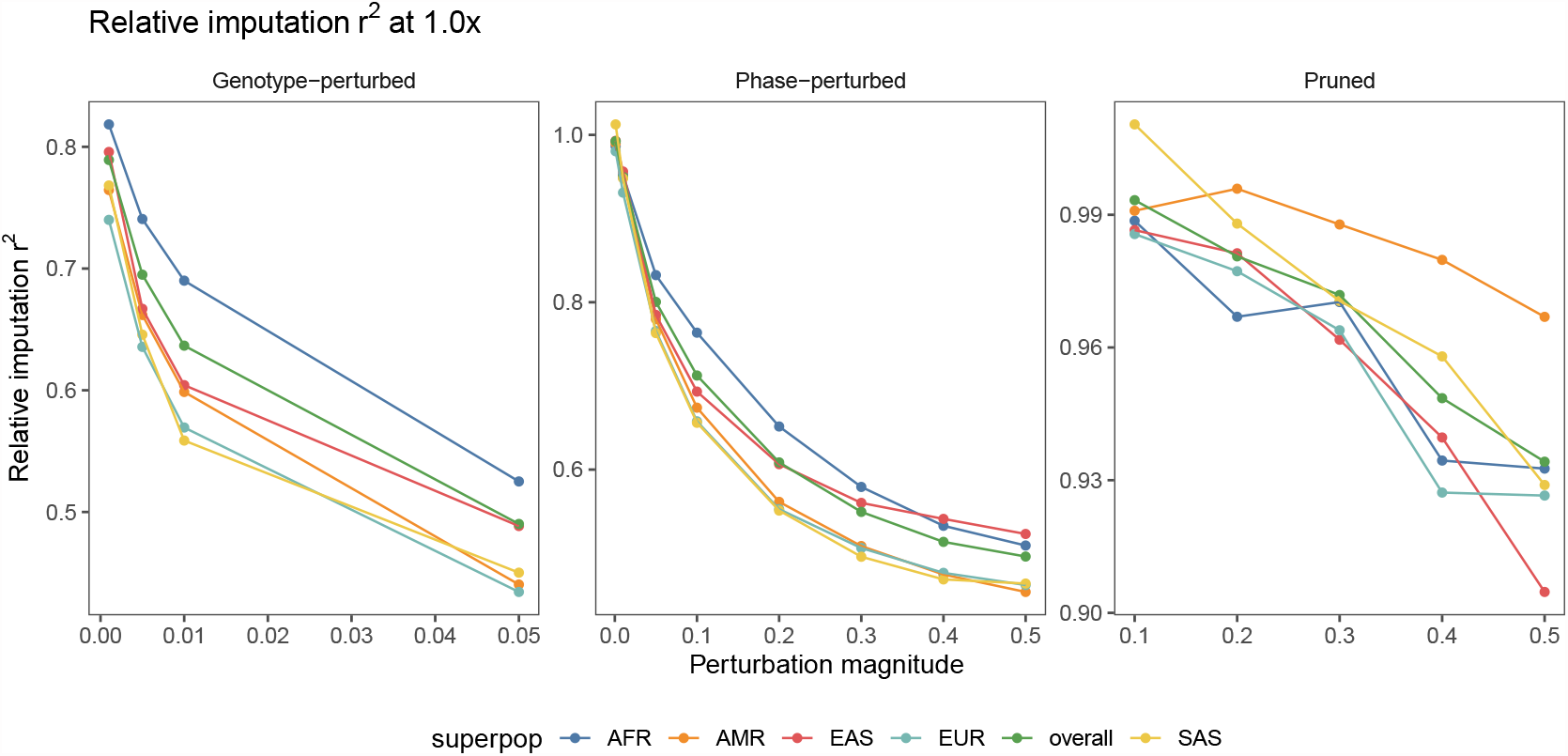
Relative (proportional) imputation *r*^2^ to the values from the unperturbed panel by perturbation magnitude, for samples at a sequencing coverage of 1.0x. Each color denotes a superpopulation group in the cohort and each pane illustrates one of the three perturbation modalities.

Three of these cases occur in the minimally phase-perturbed panel and none at higher levels of phase perturbations, leading us to hypothesize that the low level of phase error introduced was not enough to significantly attenuate overall imputation accuracy for these particular populations. The remaining cases occur in the pruned datasets at magnitudes of 0.1 and 0.3.

### Filtering low-confidence genotype calls from the haplotype reference panel

A primary conclusion of the perturbation experiments above is that imputation accuracy is sensitive to genotype error in the haplotype reference panel (Figure 1). A prediction that derives from this observation is that if low genotype quality sites can be removed from a haplotype reference panel, imputation accuracy should improve. To test this prediction, we removed all sites from chromosome 20 in the NYGC panel that had a mean genotype quality score (GQ) less than or equal to 75. GQ is a Phred-scaled measure of the confidence of the genotype call conditional on the available sequencing data for a variant, and in the case of the NYGC panel it was calculated by `GATK HaplotypeCaller` (Poplin et al., 2018).

This filter resulted in the removal of 138,241 sites, or about 9.9% of the original 1,391,020 sites. We then imputed the 60 selected individuals downsampled to 0.5x, 1.0x, and 2.0x coverage against this GQ-filtered reference haplotype panel in a leave-one-out manner (Methods) and calculated the imputation *r*^2^. When compared to the original unfiltered panel containing 1,391,020 sites, this filtered panel demonstrates an improved *r*^2^ at all three coverages (Figure 5), and the impact is particularly pronounced at MAFs ≲ 1%. Moreover, even if we take the imputed results from the original unfiltered panel and compute imputation *r*^2^ at the sites shared with the GQ-filtered panel, we still see a marginal improvement in imputation accuracy for samples imputed to the GQ-filtered panel, indicating that removal of poor-quality sites improves imputation accuracy at the remaining set of sites.

**Figure 5:**
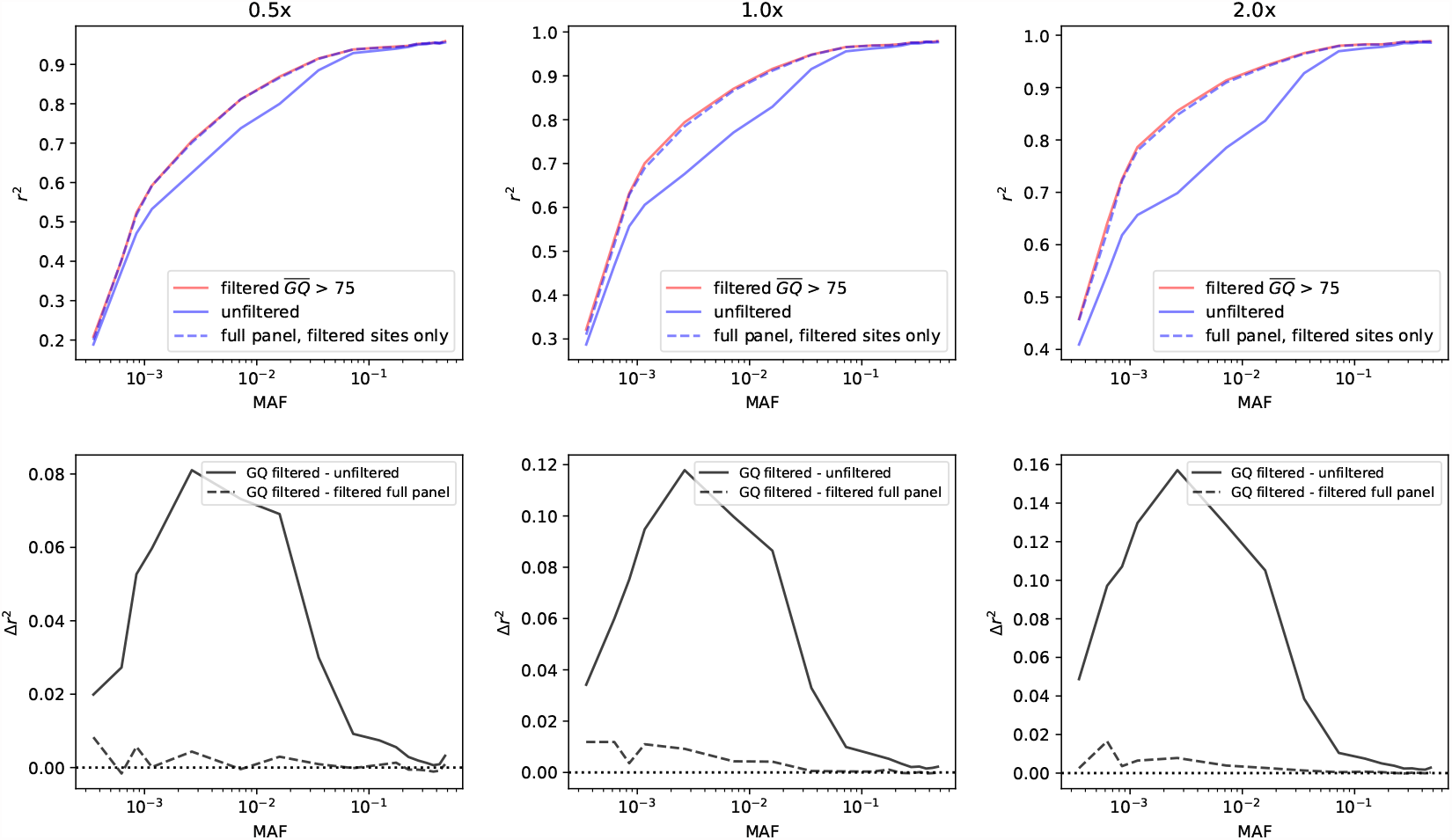
Above: imputation *r*^2^ values across MAF for the GQ filtered (red), all sites in the unfiltered panel (solid blue), and all sites imputed in the unfiltered panel but with *r*^2^ calculated only on sites intersecting with the GQ filtered panel (dashed blue) for 0.5x, 1.0x, and 2.0x coverage. Below: The imputation *r*^2^ deltas (Δ*r*^2^) across MAF for the GQ filtered panel minus the unfiltered panel (solid) and the GQ filtered panel minus the unfiltered panel but with *r*^2^ calculated only on sites intersecting with the GQ filtered panel (dashed) for 0.5x, 1.0x, and 2.0x coverage. There is also a dotted horizontal line indicating where Δ*r*^2^ equals zero.

Importantly, this filtering exercise had minimal impact on the overall panel MAF (Supplementary Figure 1), meaning the improvement in imputation *r*^2^ is a consequence of removing low quality sites from the panel rather than shifting the allele frequency spectrum.

Table 1 describes the mean genome-wide *r*^2^ (weighted by the number of variants in each allele frequency bin) resulting from imputing to the original unperturbed panel, the GQ-filtered panel, and the original panel only at sites shared with the GQ-filtered panel (*i.e*., the intersection of variants between the unfiltered and the filtered panels), at various levels of sequencing coverage. At all coverages, filtration resulted in an increase in imputation performance when compared to the original panel, ranging from an 8% increase at 0.5x to a 13% increase at 2.0x. As noted above, when restricting the comparison to the intersection of variants in the unfiltered and filtered panels, we observe a modest but nonzero improvement in *r*^2^.

In combination, these results indicate that the sites with low GQ values significantly reduce imputation performance in *cis* (i.e. the site itself has reduced accuracy), but that these low GQ sites also have a small *trans*-effect on performance (*i.e*., the low GQ sites reduce imputation accuracy at high GQ sites). Thus filtering out low GQ sites from the reference panel improves imputation in two distinct ways, both directly by removing low-quality and difficult to impute sites from the panel, but also indirectly by ameliorating low-quality linkage information when imputing high quality sites.

These empirical results support the results from simulation above and underscore the importance of genotyping accuracy in the haplotype reference panel construction process.

## Discussion

Since the first modern computational methods were developed for genotype imputation and first used in genome-wide associations in 2007, (Scott et al., 2007; Consortium et al., 2007), imputation has become a standard technique for population scale genetic studies. In the last decade and a half, sequencing costs have plummeted and algorithms have been improved, enabling the rapid imputation of genetic data to very large haplotype reference panels (Das et al., 2018). Ongoing population-scale sequencing efforts such as the UK Biobank, which recently released whole genome sequences for approximately 200 thousand individuals will lead to increasingly large and diverse reference panels, which will further spur the development and use of genotype imputation in such studies (Halldorsson et al., 2022).

As a result of the rapid development of this field, there is a rich literature in the imputation field, and dozens of papers are published each year performing comparisons of existing phasing and imputation methods, introducing new computational methods, or releasing new datasets. There are two key components in the process of genotype imputation — the haplotype reference panel and the imputation algorithm. The first represents the genetic diversity that is used in the semi-supervised learning that underlies imputation, and takes the form of a fully phased and non-missing genotype matrix. The constituent haplotypes are then used as “template” haplotypes from which a target individual’s haplotypes are modeled as imperfect mosaics. The second represents the actual implementation of the statistical model used to infer the target haplotypes, and can differ quite a bit in terms of state selection (*i.e*., the determination of the template haplotypes from which to copy) and inference methods, though the most common framework is based on the Li & Stephens hidden markov model (Li and Stephens, 2003; Das et al., 2018).

Most of the literature investigating these two factors concerns the comparison of various imputation methods or the comparison of various phasing and genotyping methods used to create reference panels. However, to the authors’ knowledge, there has yet to be a direct evaluation of the effects of phasing and genotyping errors in a reference panel on downstream imputation accuracy. Here, we aimed to fill this gap by introducing perturbations to a widely used reference panel and evaluating their effects on imputation accuracy.

For this study, we chose to use the reference panel created from the New York Genome Center’s resequencing effort of the 1000 Genomes Phase 3 dataset plus related individuals, and introduced three modes of perturbations at various magnitudes into the panel. Specifically, we introduced (1) random phase errors by flipping the phase of heterozygotes, (2) random genotyping errors by flipping the allele at genotypes, and (3) randomly pruning the reference panel. We introduced these perturbations by performing Bernoulli trials with differing rate parameters for each mode of perturbation.

Following the creation of these perturbed panels, we selected a diverse set of 60 individuals (a dozen from each superpopulation group) from the reference panel, acquired the original high-depth sequencing data for these samples, and downsampled the reads to simulate low-pass sequences at 0.5x, 1.0x, and 2.0x. We then imputed these up to the perturbed and unperturbed panels using `GLIMPSE` in a leave-one-out manner and compared the resulting imputed genotypes to the original, held out genotypes in the panel (Rubinacci et al., 2021).

We found that perturbing phase information and genotypes leads to significantly higher attenuation of imputation accuracy compared to pruning; for instance, while the maximally pruned panel with 50% of original variants removed suffered only a 6.58% decrease in mean genome-wide *r*^2^, the minimally genotype-perturbed panel with 0.01% error suffered a 21.1% reduction in the same metric for 1.0x data. Similarly, a moderately phase-perturbed panel (taking here as an example the perturbation magnitude of 0.05) resulted in a decrease of 20% in mean *r*^2^.

Notably, we found that imputation performance at high allele frequencies was surprisingly robust to perturbations, with the average imputation *r*^2^ computed at variants with MAF *≥* 10% for the 1.0x data exceeding 0.9 for all perturbation magnitudes with the exception of the 5%-genotype-perturbed panel. Conversely, the impact of phase and genotype perturbations are particularly amplified at rare variants; at the lowest allele frequency bin, 1% perturbation of genotypes decreases the mean *r*^2^ from 0.287 to 0.0433, and 5% phase perturbation decreases the *r*^2^ to 0.128. This emphasizes the importance of phasing and genotyping quality at rare variation, while placing an upper bound on the imputation accuracy decrease engendered by perturbations for common variants.

We then examined the effect of sequencing coverage of the input data, and found that the *rate of decrease* (or slope) of relative performance (computed by dividing the mean *r*^2^ from a perturbed panel to that from the unperturbed panel) with increasing perturbation magnitude decreases with increasing coverage. In other words, the negative effects of increasing perturbation magnitudes are less pronounced in higher coverage data, suggesting that the extra information from higher coverage samples ameliorates the performance deterioration induced by perturbations.

The switch error rate of a phased sample, which is a standard performance metric in the evaluation of phasing quality, was then computed on the set of phase-perturbed panels in order to place the magnitude of the random phase perturbations into a more familiar context. We found that the relationship between the magnitude of phase perturbations and induced SERs is nonlinear, and that the induced SERs ranged from 0.2% at a perturbation magnitude of 0.001 to 50% at a magnitude of 0.5. Realistically, modern phasing methods applied to high coverage whole genome sequences attain SERs of well under 1%; in this regime, the attenuation induced by switch errors is small across the allele frequency spectrum, indicating that modern reference panels likely are not missing out on significant performance gains that would be possible with better phasing algorithms.

As noted above, our simulation-based perturbation results suggest imputation accuracy is particularly sensitive to genotype accuracy in the reference panel. To determine if we could recapitulate this result in real-world data, we filtered the NYGC panel using each site’s genotype quality score (GQ), which serves as a proxy for genotype accuracy. We found that filtering out just under 10% of the sites in the panel increased genome-wide imputation accuracy, with an increase in aggregate *r*^2^ of 8 − 13% depending on coverage, and substantially more among rare alleles. This result mirrors our genotype perturbation results, where small amounts of genotype error in the reference panel rapidly decrease imputation accuracy at rare alleles, but have a more attenuated impact on common alleles. This result also suggests that our synthetic perturbations reliably predict the impact of potential filtering techniques when applied to real-world data sets. Moreover, we determined that filtration improves performance primarily by removing low-quality sites that are difficult to impute, but that it also has a small indirect benefit on the remaining high-quality sites, presumably through removing low-quality linkage information.

However, as with any other study, there are limitations. For instance, the method of perturbation we employed (*i.e*., random draws from a Bernoulli distribution) is simple, and does not model known trends such as the fact that switch errors and genotyping errors in practice are correlated with allele frequency. For future work, it may be useful to attempt to introduce perturbations in a more realistic manner that closer reflects empirical error profiles observed in real data.

Nevertheless, our results demonstrate that imputation with contemporary low-pass whole genome sequence to large haplotype reference panels is remarkably robust to realistic amounts of phase errors as well as to lower panel densities, while being sensitive to genotype error.

As increasingly large biobanks are sequenced, and as reference panels are increasingly available for agricultural species and other taxa, we anticipate that the use of genotype imputation will continue to grow, and that the sophistication of methods used for both reference panel generation and imputation itself will increase commensurately. Indeed, the utility of genotype imputation is not confined to human genomics, but has recently grown in popularity in the field of agricultural genomics, where the ability to impute unobserved genotypes has enabled more precise breeding strategies (Snelling et al., 2020; Jensen et al., 2020; Ding et al., 2023).

As genotype imputation becomes more commonplace across all applications, maintaining high quality reference panels will be essential. This study represents a comprehensive investigation into the effects of perturbations to reference panels on downstream imputation accuracy, and our results underscore the crucial role of reference panel quality for genotype imputation.

## Methods

### Sequence data

To create simulated low-pass sequencing data for the individuals under analysis, we downloaded the CRAMs made available by the NYGC from a public S3 bucket located at s3://1000genomes/ 1000G_2504_high_coverage/. We then collated the reads (to ensure pairs of reads are properly grouped) using `samtools collate` and reverted the alignments to FASTQ files using `samtools fastq` with the `-OT RG,BC` flags (Danecek et al., 2021). We then downsampled the resulting FASTQs to the equivalent of 0.5x, 1.0x, and 2.0x using `seqtk` (https://github.com/lh3/seqtk) and aligned all reads to GRCh38. Following that, we extracted only reads aligning to chromosome 20 using `samtools view` and again reverted the alignments to FASTQ using `samtools collate` and `samtools fastq`. Realignment was performed for each sample and pileups generated using `samtools mpileup`, from which genotype likelihoods were generated using the model described below. The resulting likelihoods were then used as input into a hardware-accelerated version of `GLIMPSE v1.1.1` for imputation to a filtered version of the NYGC dataset. Before perturbing the reference panel, we removed all singletons and filtered to biallelic SNPs only using `bcftools`.

### Reference panel

The phased, non-missing reference panel used in this study (the “NYGC dataset”) can be found at the following URL: http://ftp.1000genomes.ebi.ac.uk/vol1/ftp/data_collections/1000G_2504_high_coverage/working/20201028_3202_phased/. The GQ values used for filtering were derived from the original raw joint called VCFs provided at the following URL: http://ftp.1000genomes.ebi.ac.uk/vol1/ftp/data_collections/1000G_2504_high_coverage/working/20201028_3202_raw_GT_with_annot/.

### Genotype likelihoods

`GLIMPSE` takes as input genotype likelihoods (GLs) calculated at all sites in a reference panel. Although there exist a number of tools such as `bcftools` that allow the calculation of such GLs at a set of specified sites, there are a number of complications (such as `bcftools` not properly calculating GLs at indels) which led us to opt for a custom implementation of a basic GL method in a custom tool (Rubinacci et al., 2021).

At a biallelic SNP in the panel with reference and alternate alleles *a*_*r*_, *a*_*a*_ and *m* covering reads, we calculate the likelihood of a genotype *G*_*j*_ *∈* 0, 1, 2 as 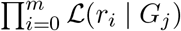 where

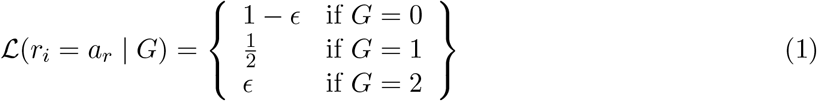

and

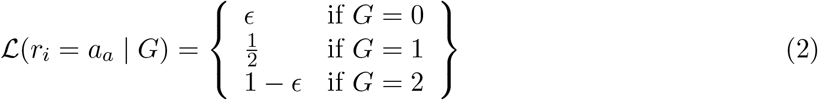

for each read *r*_*i*_, and where *ϵ* models sequencing error rate, which we set to a constant of 0.001. We then transform the resulting likelihoods into the Phred scale for input to `GLIMPSE`. For sites without reads and non-SNP sites like indels, we do not report a likelihood, which is treated by `GLIMPSE` as representing a uniform likelihood distribution.

### Imputation *r*^2^

To evaluate imputation accuracy, we computed the squared Pearson correlation coefficient (*r*^2^) between the imputed best-guess (*i.e*., the genotype with the max(GP) where GP is a three-tuple with posterior probabilities for the three possible genotypes 0*/*0, 0*/*1, 1*/*1) and the “truth” genotypes for the same individuals from the original unperturbed NYGC reference panel at each variant. As mentioned previously, this metric is useful because it has a direct interpretation in terms of statistical power and sample size in association tests for binary traits. Specifically, the statistical power of a test with a sample size of *N* at an imputed variant with imputation *r*^2^ = *a* is approximately equal to the power of a test with sample size *aN* when the genotypes at that variant are known and accurate. Thus, the average imputation *r*^2^ genome-wide can be thought of as measuring the average effective *sample size reduction* suffered when using imputed genotypes vs. the ideal case in which all variants are known (Das et al., 2018; Pritchard and Przeworski, 2001). Since this metric is undefined if either the vector of imputed and true genotypes have zero covariance, which is often the case at rare variants, we report an *aggregate r*^2^, which is the squared correlation coefficient between the vector of all genotypes for imputed variants aggregated in a minor allele frequency bin and the same for the truth genotypes (Hoffmann and Witte, 2015; Rubinacci et al., 2021; 1000 Genomes Project Consortium, 2015; Marchini, 2019).

Note that while this aggregation method circumvents the problem of invariant genotype vectors, it is *not* mathematically equivalent to the mean of the imputation *r*^2^ in a given AF bin; thus, this metric should be evaluated on its own terms and only as a heuristic or estimator of the true imputation *r*^2^ statistic as originally defined.

We used the `GLIMPSE2_concordance` tool to calculate the aggregate imputation *r*^2^ in each allele frequency bin for each cohort, where the bins have boundaries set at 0.0001, 0.0005, 0.00075, 0.001, 0.00125, 0.005, 0.01, 0.0125, 0.05, 0.1, 0.15, 0.2, 0.25, 0.3, 0.35, 0.4, 0.45, 0.5. For each level of coverage and perturbation magnitude, we computed the aggregate *r*^2^ for the overall cohort as well as for each superpopulation subset, in all cases using the genotypes in the unperturbed reference panel as the comparison “truth” set.

To calculate an average *r*^2^ across some frequency range, we multiplied the aggregate *r*^2^ in a bin by the number of variants in that bin, summed these values across the bins of interest, and divided the sum by the total number of variants in the bins of interest; that is, we calculate the average *r*^2^ from bins *i* to *j* as

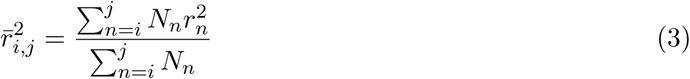

where *N*_*n*_ is the number of variants in bin *n*.

## Author contributions

J.H.L., L.F., and C.A.B. were involved in study design and conceptualization. J.H.L., A.L., C.A.B., W.P., G.M.B., M.A., M.J.S.G., and L.F. were involved in statistical and computational analysisJ.H.L. and L.F. wrote the original manuscript draft.

## Footnotes

1 N.B.: all discussion of allele frequencies in this manuscript are with respect to the frequency spectrum in the full, unperturbed reference panel

