## Supplementary Materials for "The effects of reference panel perturbations on the accuracy of genotype imputation" (from another version of this preprint: version 1, 2023-08-10)

### 1 Figures

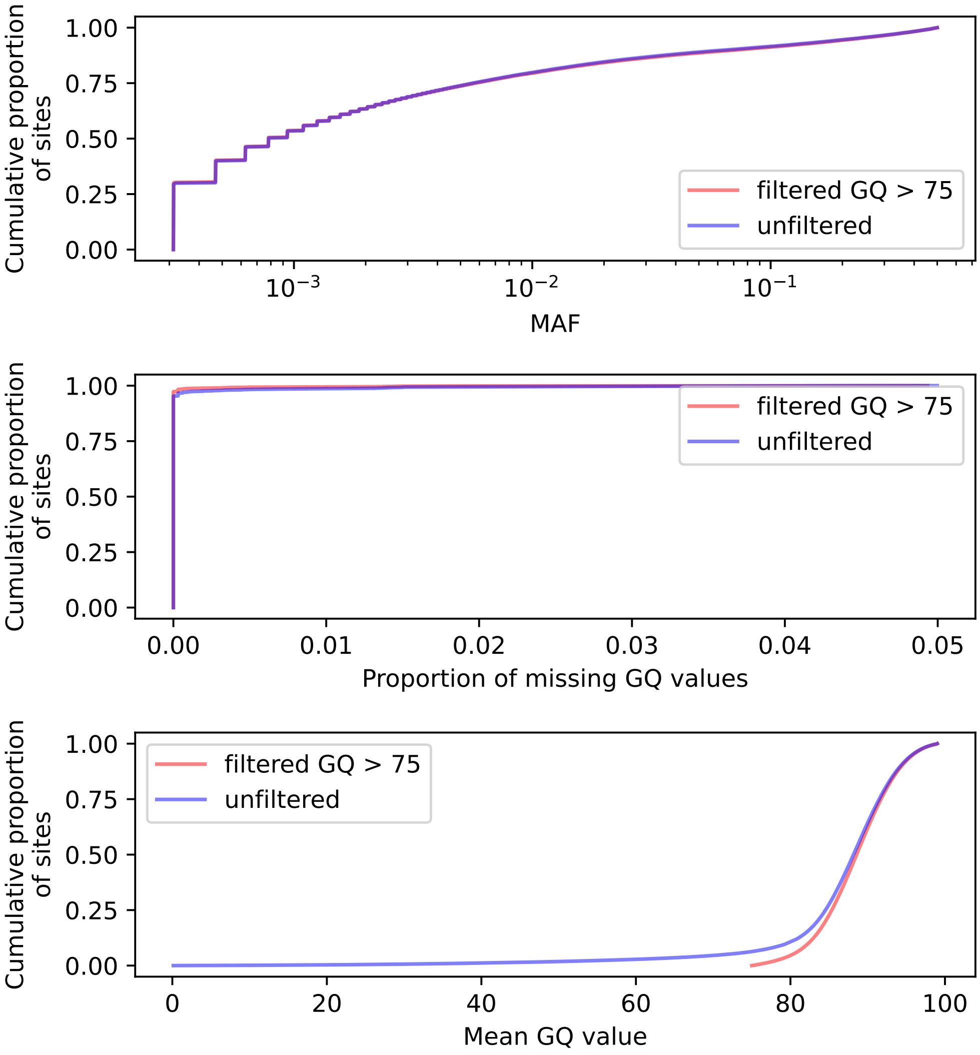


**Supplementary Figure 1:** The impact of removing sites with mean GQ< 75 on 1KGP3 panel statistics including minor allele frequency (MAF), missing GQ values, and on GQ itself. Note that "missing GQ values" exist due to the fact that we took the GQ values from the raw joint-called genotypes available for the panel, since the phased callset did not contain this information.

#### 1.1 Tables

| Superpopulation group | Coverage | Magnitude | Relative $r^2$ | Perturbation |
|---|---|---|---|---|
| AMR | 0.5x | 0.001 | 1.005896 | phase |
| AMR | 2.0x | 0.300 | 1.001743 | prune |
| EAS | 0.5x | 0.001 | 1.003715 | phase |
| SAS | 1.0x | 0.001 | 1.012525 | phase |
| SAS | 1.0x | 0.100 | 1.010436 | prune |

**Supplementary Table 1:** Table describing the datasets in which *greater* mean imputation $r^2$ is achieved compared to the unperturbed panel. Here, relative $r^2$ is the mean genome-wide $r^2$ for a dataset divided by the same metric from the unperturbed panel.
